# Characterization and Genome Insights of Two Novel Plastic-Degrading *Acinetobacter* Species

**DOI:** 10.1101/2025.07.16.665163

**Authors:** Mengli Xia, Yuandong Zhao, Yiran Ma, Bo Wu, Guoquan Hu, Yanwei Wang, Mingxiong He

## Abstract

The ubiquitous presence of *Acinetobacter* species in the environment holds significant biotechnology potential, particularly in the degradation of various pollutants. In this study, we present two plastic-degrading bacteria, strains CAAS 2-6^T^ and CAAS 2-13^T^, isolated from landfill leachate and a strawberry farmland, respectively. Strain CAAS 2-6^T^ showed the highest *16S rRNA* sequence similarity (97.7%) with *Acinetobacter gerneri* DSM 14967^T^, while strain CAAS 2-13^T^ was most closely related (99.6%) to *Acinetobacter gerneri* PS-1^T^. Phylogenetic analysis of *16S rRNA* and *gyrB*-*rpoB* genes placed both strains on distinct branches. Genomic comparisons revealed the highest digital DNA–DNA hybridization (dDDH) / average nucleotide identity (ANI) values for CAAS 2-6^T^ with *Acinetobacter indicus* CIP 110367^T^ (22.7%/77.0%) and for CAAS 2-13^T^ with *Acinetobacter gerneri* PS-1^T^ (66.5%/96.0%). Based on phenotypic and chemotaxonomic data, we propose strain CAAS 2-6^T^ as the novel species *Acinetobacter lentus* sp. nov. (type strain CAAS 2-6^T^ = GDMCC 1.3951^T^ = JCM 36321^T^ = KCTC 8156^T^), and strain CAAS 2-13^T^ as the novel subspecies *Acinetobacter kanungonis* subsp. *fragariae* subsp. nov. (type strain CAAS 2-13^T^ = GDMCC 1.3956^T^ = JCM 36322^T^ = KCTC 8157^T^). Both strains utilized PLA, PBSA, PBS, PBAT, and PBT as sole carbon sources, with CAAS 2-6^T^ exhibiting superior growth on PBSA, PBS, and PBAT. Genomic annotation identified genes encoding plastic-degrading enzymes multicopper oxidase AbMCO and alkane hydroxylase AlkB. Enzymatic depolymerization assays confirmed *in vitro* production of plastic monomers. These findings expand the known metabolic capabilities of *Acinetobacter* and provide promising new candidates for plastic bioremediation.

**IMPORTANCE:** *Acinetobacter* spp. demonstrate significant potential for plastic bioremediation. We characterize two novel strains: CAAS 2-6^T^ (*A. lentus* sp. nov.) and CAAS 2-13^T^ (*A. kanungonis* subsp. *fragariae* subsp. nov.), capable of degrading five major polyesters (PLA, PBAT, PBT, PBS, PBSA) as sole carbon sources. Genomic and enzymatic analyses identified multicopper oxidase (AbMCO) and alkane hydroxylase (AlkB) as key depolymerase. This work: (i) Expands plastic-degrading *Acinetobacter* diversity; (ii) Provides novel biocatalysts and genomic resources; (iii) Delivers engineerable microbial chassis.

## INTRODUCTION

The genus *Acinetobacter*, introduced by Brisou and Prévot (1), belongs to the family *Moraxellaceae*, order *Moraxellales*, and class *Gammaproteobacteria*. This taxonomically diverse and ubiquitous genus comprises 87 species with validly published names (https://lpsn.dsmz.de/genus/acinetobacter; last accessed June 17, 2025). *Acinetobacter* spp. are Gram-negative, aerobic, catalase-positive and oxidase-negative, with genomic G+C content of 34.9–47.0% (2). They inhabit diverse environments, including soil (3, 4, 5, 6, 7, 8), water ecosystems (2, 9, 10, 11, 12), plants (13, 14), animals (15, 16, 17, 18, 19, 20), and hospital environments (21, 22, 23) (apps.szu.cz/anemec/Classification.pdf), reflecting broad metabolic versatility (24).

Several *Acinetobacter* species exhibit significant biotechnological potential through the degradation of diverse pollutants, including plastics, hydrocarbons, pesticides, and plasticizers. *Acinetobacter* spp. metabolizing pesticides (methyl parathion (25), methamidophos (26), fenamidophos (27), bensulfuron-methyl (28)), and diesel alkanes (e.g., *A. baumannii* degrading 58.1% in 10 days (29); *A. vivianii* KJ-1 utilizing diesel as sole carbon source (30); *A. junii* degrading 57.5% in 45 days (31); *A. venetianus* RAG-1 participates in long-chain alkane degradation via alcohol dehydrogenases (ADHs) (32); *A. haemolyticus* JS-1 degrade 70–80% of crude oil at concentrations of 10–20 g/L within 15 days at 30°C (33)). Strains degrading phthalate esters (e.g., *Acinetobacter* sp. SN13 on di(2-ethylhexyl) phthalate (DEHP) (34); strain LMB-5 on multiple phthalates (35)). Plastic-degrading strains such as *A. gerneri* P7 (polyurethane via 66-kDa esterase (36)), *A. pittii* IRN19 (untreated LDPE (37)), *Acinetobacter* sp. AnTc-1 (polystyrene (38)), and gut-derived *Acinetobacter* sp. BIT-H3 (rubber (39)). Enzymatic drivers like AlkB (alkane degradation (30, 40)) and AbMCO (polyethylene degradation (41)). The novel species A. suaedae sp. nov. preferentially degrades phenolic acids (42), further highlighting metabolic versatility. Despite their metabolic diversity, only a limited number of *Acinetobacter* species have been definitively associated with plastic degradation, and the taxonomic relationship between degradation capacity and phylogenetic position remains unclear. This knowledge gap underscores the need for polyphasic characterization of novel plastic-degrading isolates.

Improper plastic disposal threatens ecosystems and health (43). Microbial degradation offers a promising remediation strategy (44, 45), yet key resources are underexplored. Landfills store 21–42% of global plastic waste (46), while agricultural soils accumulate plastic film residues (47)——both likely harbor specialized plastic-degrading microbes. Despite this potential, *Acinetobacter* strains capable of metabolizing multiple plastics remain scarce, and their enzymatic machinery is poorly characterized.

Here, we isolated two novel strains, CAAS 2-6^T^ and CAAS 2-13^T^, from environmental samples. Both utilize multiple plastics as sole carbon sources. To resolve their taxonomy, we conducted phylogenetic analyses (16S rRNA, gyrB-rpoB), genome-based digital DNA-DNA hybridization (dDDH) and average nucleotide identity (ANI). We further mined their genomes for he multicopper oxidase gene *abMCO* and the alkane 1-monooxygenase gene *alkB* homologs, heterologously expressed these in *Escherichia coli* BL21 (DE3), and validated enzymatic depolymerization of biodegradable polymers. Our work establishes CAAS 2-6^T^ and CAAS 2-13^T^ as the novel species and a novel subspecies *Acinetobacter lentus* sp. nov. and subspecies *Acinetobacter kanungonis* subsp. *fragariae* subsp. nov., respectively. This represents the first systematic validation of multi-plastic degradation in *Acinetobacter* via integrated taxonomy and enzymology.

## MATERIALS AND METHODS

### Reagents and culture media

This study utilized a plastic film made of Polylactic acid (PLA) donated by Sichuan Shennong Chinese Medicine Agricultural Research Institute Group Co., Ltd., and six types of plastics were utilized. Among them, PLA was sourced from NatureWorks LLC in the United States. Polybutylene adipate terephthalate (PBAT) and Polybutylene succinate-adipic acid (PBSA) were produced by Xinjiang Lanshantunhe Science and Technology Co., Ltd in China. Polyethylene terephthalate (PET) came from DuPont de Nemours, Inc. in the United States. Polybutylene terephthalate (PBT) was manufactured by Chang Chun Plastics and Chemical Industry Co., Ltd. in China, and Polybutylene succinate (PBS) was supplied by PTT MCC Biochem Co., Ltd. in Thailand. Other chemical reagents were purchased from Sinopharm Chemical Reagent Co., Ltd. and Shanghai Macklin Biochemical Co.,Ltd. in China. KOD One polymerase was purchased from Takara Biotechnology Co., Ltd. in Japan. Mineral salt medium (MSM) consisted the following (g/L): 7.0 K_2_HP_4_O, 2.0 KH_2_PO_4_, 1.0 NH_4_NO_3_, 0.1 MgSO_4_·7H_2_O, 0.01 FeSO_4_·7H_2_O, 0.001 ZnSO_4_·7H_2_O, 0.0002 MnSO_4_·6H_2_O and 0.0001 CuSO_4_·7H_2_O (pH 7.5) (48, 49, 50). Add 1.0 g/L yeast extract as a growth factor if necessary. Luria-Bertani medium (LB) consisted of (g/L): 10.0 Tryptone, 5.0 yeast extract and 5.0 NaCl. The solid medium was supplemented with 15 g/L agar and sterilized at 121°C for 20 minutes.

### Strain isolation and molecular identification

This study conducted sampling from various environments in Chengdu, Sichuan Province, China. 2 g of each sample was enriched in five rounds, with each round lasting for 7 days, in 25 mL of MSM medium supplemented with PLA film. The PLA film was sterilized by soaking in 75% ethanol for 10 minutes and then air-dried under UV light. Enrichment and subculturing were performed by transferring 2 mL of the pelagic microbes and PLA film from the previous culture to fresh 25 mL of MSM. Following the completion of the fifth enrichment round, *Acinetobacter* species were isolated and purified from the culture fluid. The standard dilution plating method was employed to spread the enriched culture onto LB agar plates (51). After incubating at 30°C for 72 hours, individual colonies were randomly selected and repeatedly streaked on LB agar plates to obtain pure cultures. The selected purified isolates were cryopreserved in a suspension containing 20% (v/v) glycerol at −80°C. For phylogenetic analysis of the *16S rRNA* gene, PCR amplification and sequencing were carried out following a previously described method (52). Subsequent BLAST analysis was performed for the *16S rRNA* gene sequences via the EzBioCloud web-service (53).

### Phenotypic and chemotaxonomic characterization

Colony morphology was examined on LB agar after 48-h incubation at 37°C. Cell motility was assessed through stab inoculation in semi-solid medium containing 0.5% (w/v) agar. Gram reaction was determined according to the instructions of the commercial kit (G1060; Beijing Solaibao Technology Co., Ltd in China). Anaerobic growth capability was evaluated in Mitsubishi™ AnaeroPouch-Bag systems during a 7-day incubation at 30°C. Scanning electron microscopy (SU8010; Hitachi, Ltd. in Japan) analysis of mid-log phase cells cultured in LB broth at 30°C revealed cellular morphology. Growth parameters were determined by measuring optical density at 600 nm (L5S; INESA (Group) Co., Ltd in China) in LB medium after 48-h incubation under controlled conditions: thermal gradients (4, 8, 16, 20, 25, 30, 37, 40, 45, 50 and 60°C), pH adaptability (4.0–11.0 in 0.5-unit increments with biological buffers), and halotolerance (0–10% (w/v) NaCl at 1% intervals). All physiological tests were conducted in biological triplicates. Catalase activity was confirmed through bubble formation in 3% (v/v) hydrogen peroxide, with oxidase activity detected using 1% (w/v) tetramethyl-ρ-phenylenediamine. Acid production from carbohydrates was analyzed using the API 50CH system (bioMérieux), while other biochemical and enzymatic tests were performed with the API 20E, API 20NE and API ZYM kit (bioMérieux) following manufacturer protocols.

For the determination of the whole-cell fatty acid composition, strains were cultivated on LB agar plate at 30°C for 48 hours. The fatty acids were then extracted and analyzed according to the recommendations of the Microbial Identification System (MIDI), a commercial identification system (54). The whole fatty acid composition was determined using an Agilent 6890N gas chromatograph. For the analysis of lipids, cells were cultured in LB culture until the mid-exponential phase on a rotary shaker at 30°C. Polar lipids were extracted from 200 mg lyophilized biomass using the Bligh and Dyer method (55), followed by two-dimensional thin-layer chromatography (2D-TLC) separation on silica gel GF254 plates (100*×*100 mm; Qingdao Marine Chemical Factory Branch in China) under chromatographic conditions established according to Minnikin et al. (56, 57, 58). Definitive lipid profiling and validation were subsequently performed by the China Agricultural Culture Collection Center (ACCC) through standardized microbial chemotaxonomic protocols.

### Phylogenetic analyses based on *16S rRNA* and *gyrB*-*rpoB* gene sequences

The 16S rRNA gene sequences of strains CAAS 2-6^T^ and CAAS 2-13^T^, along with 99 typical strains of the genus Acinetobacter available in the NCBI database, were aligned using CLUSTAL X 2.0 software (59). To further determine the taxonomic position of the strains, a multilocus sequence analysis (MLSA) was performed on the housekeeping genes, DNA gyrase subunit B (*gyrB*) and RNA polymerase β-subunit (*rpoB*), from the whole-genome sequences. The aligned sequences were used in the program MEGA 11.0 (60) to construct the neighbour-joining (NJ) (61) and maximum-likelihood (ML) (62) phylogenetic trees using the Maximum Composite Likelihood and Tamura-Nei model, with bootstrap values calculated from 1000 replications.

### Whole genome sequencing and analysis

The draft genome was completed by Shanghai Pasino Biotechnology Co., Ltd. in China through the Whole Genome Shotgun strategy. Libraries with different insert sizes were constructed and sequenced via second-generation Next-Generation Sequencing technology on the Illumina NovaSeq platform using Paired-end sequencing. To maintain data integrity, AdapterRemoval (63) eliminated splicing contaminants at the 3’ end, while SOAPec (64) performed quality corrections. The reads were subsequently de novo processed and assembled using A5-MiSeq version 20160825 (65) along with SPAdes v3.12.0 (66) (http://cab.spbu.ru/files/release3.12.0/manual.html). Based on the results of SPAdes, base correction was performed using Pilon (67) software v1.18. GeneMarkS v4.32, Barrnap v0.9, tRNAscan-SE v1.3.1, and CRISPRCasFinder v4.2.20 were used for predicting and annotating open reading frames, rRNA, tRNA, and CRISPRs (68, 69, 70, 71). The encoded protein sequences underwent comparison with those in the NCBI NR, eggNOG (evolutionary genealogy of genes: Non-supervised Orthologous Groups), KEGG (Kyoto Encyclopedia of Genes and Genomes), Swiss-Prot, and GO (Gene Ontology) databases via diamond blastp. The genome circos map was constructed by CGView (72).

A preliminary phylogenetic assessment of strains CAAS 2-6^T^ and CAAS 2-13^T^ was conducted using the Type Strain Genome Server (TYGS) tool (https://tygs.dsmz.de, accessed20October2023) for genomic clustering analysis (73). Subsequently, dDDH and ANI values between these strains and closely related *Acinetobacter* strains were calculated via the EzBioCloud web server (https://www.ezbiocloud.net/tools/ani) (53) and the Genome-to-Genome Distance Calculator (GGDC) v3.0 using the recommended Formula 2 (http://ggdc.dsmz.de) (74). Additionally, genome-wide relatedness indices were evaluated using BLAST-based ANI (ANIb) (75) and OrthoANI (76). Finally, genomic collinearity analysis of chromosomes, genes, and their start/end positions among strains CAAS 2-6^T^, CAAS 2-13^T^, and *A. kanungonis* PS-1^T^ was performed using TBTools-II v2.225 (77).

### Growth studies in modified minimal salt media supplemented with plastics

Cells were cultured in LB medium at 30°C to mid-log phase (OD_600_*≈*0.6), harvested by centrifugation (4,000 *×*g, 3 min, 4°C), washed thrice with sterile 0.85% (w/v) NaCl, and resuspended in modified mineral salt medium (MSM, 1.0 g/L yeast extract). For growth utilization assays, saline-washed cells (OD_600_ adjusted to 0.8 *±* 0.05) were inoculated at 10% (v/v) into 9 mL of MSM with 1.0 g/L yeast extract supplemented with 0.2 g/L plastic powder (<100 µm particle size) as sole carbon source. Cultures were incubated at 30°C with shaking (180 rpm) for 7 days. Biomass accumulation was monitored at OD_600_, followed by HPLC analysis of culture supernatants for monomer quantification.

### HPLC analyze

The liquid samples were centrifuged at 12,000 *×*g for 10 min and filtered through 0.22 µm membranes into HPLC vials. Analytical protocols were tailored to monomer types by selecting distinct instruments, columns, and programs. For nonpolar or weakly polar compounds, a Shimadzu LC-16 HPLC system equipped with an Agilent HC-C18 column (4.6 *×* 250 mm, 5 µm particle size) was employed. The mobile phase consisted of 0.1% (v/v) formic acid and acetonitrile under gradient elution: 70% formic acid/30% acetonitrile (0–10 min), linear gradient to 100% acetonitrile (10–20 min), and 100% acetonitrile (20–30 min). The injection volume was 20 µL, column temperature maintained at 30°C, and flow rate set to 1.0 mL/min, with effluent monitored at 254 nm using a diode array detector (DAD). For polar compounds, an Agilent 1200 HPLC system with an Aminex HPX-87H column (300 *×* 7.6 mm) was utilized. Isocratic elution was performed over 30 min using 50 mM sulfuric acid as the mobile phase. The injection volume was 20 µL, column temperature held at 35°C, and flow rate adjusted to 0.6 mL/min, with effluent detected by a refractive index detector (RID). Calibration curves were constructed using standard solutions of terephthalic acid (TPA), lactic acid (LA), 1,4-butanediol (BDO), adipic acid (AA), and succinic acid (SA) at concentrations of 0.05, 0.1, 0.25, 0.5, 1.0, 2.5, and 5.0 g/L. Triplicate injections were performed for each concentration, and linear regression analyses yielded correlation coefficients R^2^ >0.999.

### Gene cloning, protein purification and enzymatic degradation of plastic powder

Gene sequences encoding putative multicopper oxidase AbMCO (EC 1.3.3.5) and alkane hydroxylase AlkB 1.14.15.3) homologs were selected for molecular cloning and heterologous expression. Target genes were PCR-amplified using gene-specific primers (Table S2; synthesized by Sangon Biotech, Shanghai) and subsequently cloned into the pET28a(+) expression vector via Gibson assembly. The recombinant plasmids were transformed into *Escherichia coli* BL21(DE3) cells for polyhistidine-tagged protein purification as follows:

(1) Primary seed cultures were grown overnight at 37°C with 200 rpm shaking in 5 mL LB broth supplemented with kanamycin (0.1 g/L). Secondary cultures were inoculated at 1:50 dilution in 50 mL kanamycin-supplemented LB broth (0.1 g/L) and incubated at 37°C (200 rpm) until OD_600_ reached 0.6–0.8. Protein expression was induced with 1 mM IPTG at 25°C for 10 h (200 rpm). Cells were harvested by centrifugation at 4,000 *×*g for 10 min (4°C; TGL-21M centrifuge, Luxiangyi).
(2) Cell pellets were resuspended in lysis buffer and disrupted by ultrasonication (250 W; 5 s pulse-on/5 s pulse-off cycles for 15 min total; SCIENTZ-IID, Xinzhi). Lysates were clarified by centrifugation at 20,000 *×*g for 20 min (4°C; TGear HC16R, Tiangen). Supernatants were subjected to affinity purification using Ni-NTA gravity columns (HyPur™ T 6FF; Sangon Biotech, C600791-0010) according to manufacturer protocols.
(3) Protein expression and purity were verified by SDS-PAGE. Concentrations were determined via modified Bradford assay (Sangon Biotech, C503041-1000) using bovine serum albumin (BSA) standards.
(4) A 200 µL aliquot of the nickel column-purified protein solution was mixed with 800 µL of either 50 mM sodium acetate buffer (pH 6.0, 3.88 g/L CH_3_COONa and 0.155 mL/L CH_3_COOH) or 50 mM Tris-HCl buffer (pH 7.5, 6.06 g/L Tris and 3.33 mL/L HCl) supplemented with 1 mM NADH. The reaction mixtures were incubated at 37°C with agitation (180 rpm) for 5 days. Post-incubation supernatants of the enzymatic hydrolysate were collected, filtered to sterilize, and then subjected to HPLC analysis to quantify the production of plastic monomers.

### Modeling and molecular docking

Protein structure prediction was performed using AlphaFold 3 (78) via Google Colaboratory, with the optimal model validated by Ramachandran plot analysis. Substrate molecules were sketched in ChemDraw 3D (v20.0) and exported as PDB files. Molecular docking was conducted in AutoDock Vina (79, 80) following hydrogen addition to both the predicted enzyme structure and substrate. Three-dimensional structural visualization and docking analyses were performed using PyMOL (v2.6).

Molecular docking studies were performed using oligomers composed of six monomer candidates—terephthalic acid (T), 1,4-butanediol (B), ethylene glycol (E), adipic acid (A), succinic acid (S), and lactic acid (L)—as substrates. The selected oligomers (ABT, BAB, BABTB, BSB, BT, BTB, BTBT, LLLL, MHET, MHET_4_, TBT, TBTBT) were docked against the predicted 3D structures of AlkB enzyme from strain CAAS 2-13^T^ and AbMCO1 enzyme from strain CAAS 2-6^T^. To determine the conserved domain characteristics, the protein structures of the same enzyme from different species within the same genus were also compared.

## RESULTS AND DISCUSSION

### Isolation, morphology and chemotaxonomic features

The strain CAAS 2-6^T^ was isolated from leachate of Chang’an Landfill (coordinates: 30*^◦^*39*^′^*12.74*^′′^*N, 104*^◦^*22*^′^*41.44*^′′^*E) in the Longquan Mountain Range, Chengdu, China, while strain CAAS 2-13^T^ originated from a strawberry cultivation field (coordinates: 30*^◦^*24*^′^*33.27*^′′^*N, 104*^◦^*09*^′^*05.66*^′′^*E) on the Chengdu Plain. Both strains formed circular, smooth-surfaced colonies with entire margins and convex elevation on LB agar, measuring 0.5–1.0 mm (CAAS 2-6^T^) and 1.5–2.0 mm (CAAS 2-13^T^) in diameter (Fig. S1a). The cells were Gram-stain-negative, anaerobic, and short rods with dimensions of 0.6–0.7 µm *×* 0.7–1.3 µm (CAAS 2-6^T^) and 0.5–0.6 µm *×* 0.7–1.2 µm (CAAS 2-13^T^) (Fig. **1**). Physiological characterization revealed optimal growth conditions for CAAS 2-6^T^ at 25–40°C, pH 7.0–7.5, and 0–4.0% NaCl tolerance (range: 15–40°C, pH 6.0–8.0, 0–6.0%), and for CAAS 2-13^T^ at 25–37°C, pH 5.0–7.5, and 0–3.0% NaCl (range: 15–40°C, pH 4.5–8.0, 0–5.0% NaCl), with supporting data provided in Fig. S1.

**FIG 1.**
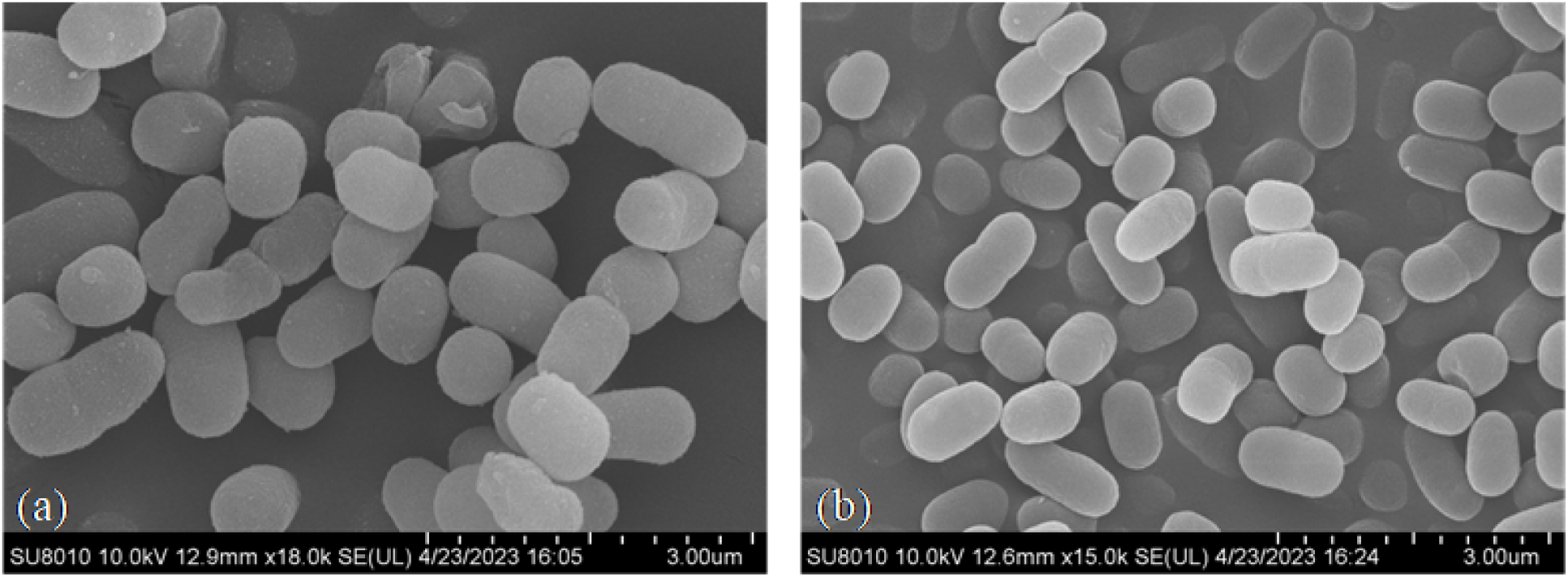
Scanning electron micrograph (SEM) of strain CAAS 2-6^T^. (a) and CAAS 2-13^T^ (b) cells showing the rod-shape with the size of approx. 0.6–0.7 µm *×* 0.7–1.3 µm and 0.5–0.6 µm *×* 0.7–1.2 µm. Bar, 3 µm.

Both CAAS 2-6^T^ and CAAS 2-13^T^ exhibit oxidase-negative and catalase-positive. Comparative phenotypic analysis of strains CAAS 2-6^T^, CAAS 2-13^T^, and *A. gerneri* KCTC 12415^T^ is presented in Table S2. API ZYM/20E/20NE/50CH profiles confirmed these strains as members of the genus *Acinetobacter*. Consistent with the genus-specific enzymatic profile, neither strain produced arginine dihydrolase, lysine decarboxylase, ornithine decarboxylase, trypsin, chymotrypsin, α-galactosidase, β-galactosidase, α-glucosidase, β-glucosidase, β-glucuronidase, N-acetyl-β-glucosaminidase, α-mannosidase, or β-fucosidase. Both strains exhibited alkaline phosphomonoesterase, acid phosphomonoesterase, esterase (C4), esterase lipase (C8), and leucine arylamidase activities. The following strain-specific characteristics were identified: CAAS 2-6^T^ exhibited gelatinase negativity coupled with atypical metabolic capabilities including acid production from multiple substrates and assimilation of mannose, sucrose, and starch - characteristics divergent from all validated *Acinetobacter* spp.; CAAS 2-13^T^ demonstrated restricted catabolic competence, showing weak acidification of sorbitol, D-mannitol, and sucrose, along with failure to assimilate D-glucose, adipic acid, or utilize various carbon sources, thereby differentiating it from other members of the genus.

The whole-cell fatty acid profiles of strains CAAS 2-6^T^, CAAS 2-13^T^ and *A. gerneri* KCTC 12415^T^ are presented in Table **1**. Strain CAAS 2-6^T^ contained Summed Feature 3 (C_16:1_ *ω*7*c* and/or C_16:1_ *ω*6*c*, 25.8%), C_16:0_ (25.7%), and C_18:1_ *ω*9*c* (10.5%) as major components, while strain CAAS 2-13^T^ showed predominant levels of Summed Feature 3 (24.1%), C_16:0_ (25.5%), and C_18:1_ *ω*9*c* (21.6%). The polar lipid profiles of strain CAAS 2-6^T^ and *A. gerneri* KCTC 12415^T^ comprised diphosphatidylglycerol (DPG), phosphatidylethanolamine (PE), phosphatidylglycerol (PG), aminolipid (AL), unidentified phosphoglycolipids (PL), aminophospholipids (APL), unknown polar lipids (L) (Fig. S2). These chemotaxonomic characteristics aligned with those of recently described *Acinetobacter* species (24, 35, 81), confirming the phylogenetic placement of both strains within the genus *Acinetobacter*.

**TABLE 1.**
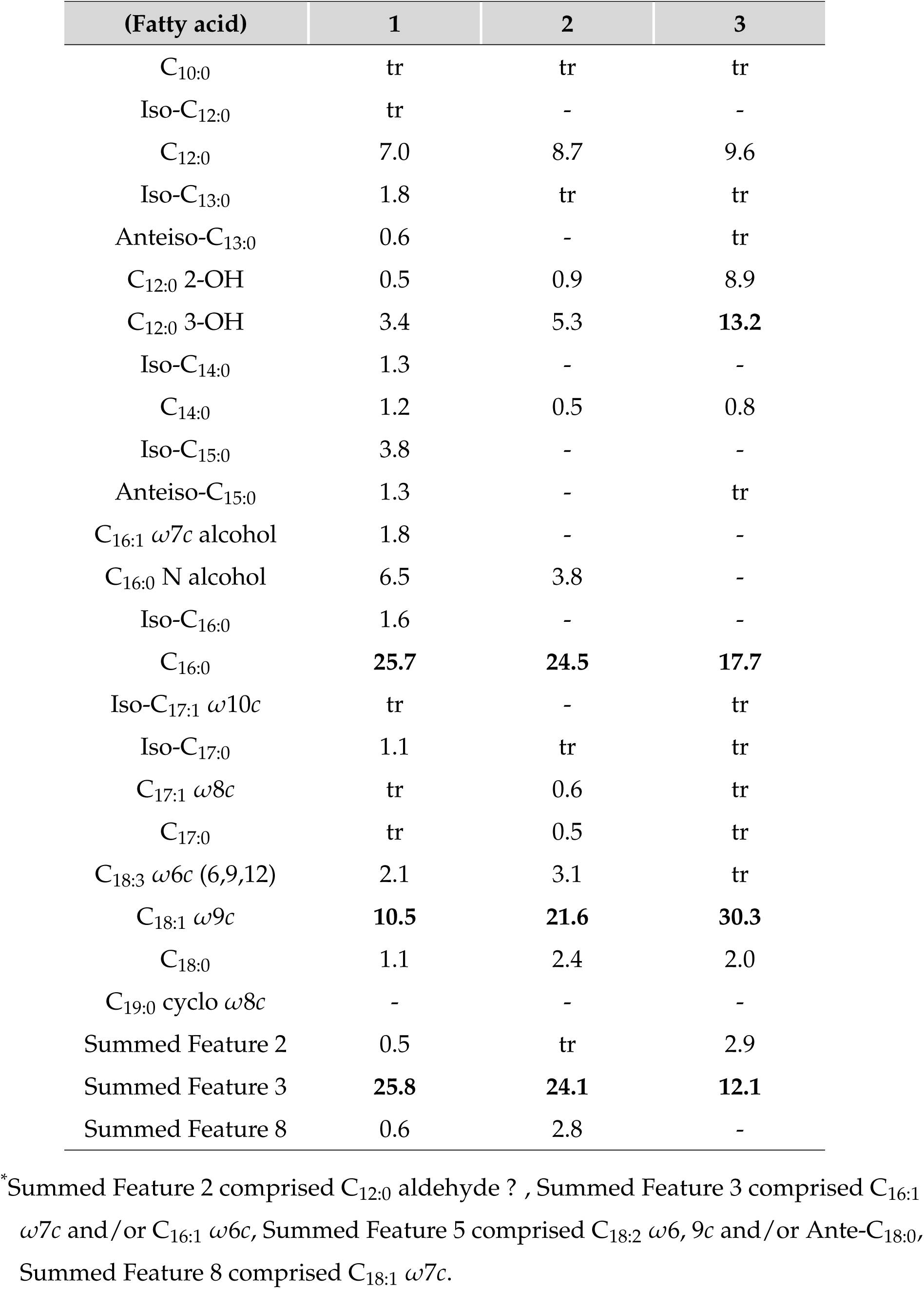
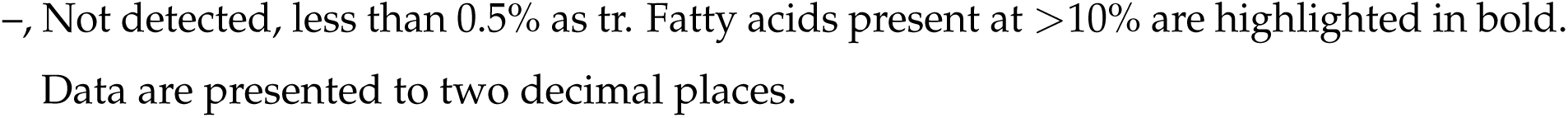
Fatty acid results of (1) strains CAAS 2-6^T^, (2) CAAS 2-13^T^, and (3) *A. gerneri* KCTC 12415^T^.

| (Fatty acid) | 1 | 2 | 3 |
| --- | --- | --- | --- |
| C<sub>10:0</sub> | tr | tr | tr |
| Iso-C<sub>12:0</sub> | tr | - | - |
| C<sub>12:0</sub> | 7.0 | 8.7 | 9.6 |
| Iso-C<sub>13:0</sub> | 1.8 | tr | tr |
| Anteiso-C<sub>13:0</sub> | 0.6 | - | tr |
| C<sub>12:0</sub> 2-OH | 0.5 | 0.9 | 8.9 |
| C<sub>12:0</sub> 3-OH | 3.4 | 5.3 | **13.2** |
| Iso-C<sub>14:0</sub> | 1.3 | - | - |
| C<sub>14:0</sub> | 1.2 | 0.5 | 0.8 |
| Iso-C<sub>15:0</sub> | 3.8 | - | - |
| Anteiso-C<sub>15:0</sub> | 1.3 | - | tr |
| C<sub>16:1</sub> ω7*c* alcohol | 1.8 | - | - |
| C<sub>16:0</sub> N alcohol | 6.5 | 3.8 | - |
| Iso-C<sub>16:0</sub> | 1.6 | - | - |
| C<sub>16:0</sub> | **25.7** | **24.5** | **17.7** |
| Iso-C<sub>17:1</sub> ω10*c* | tr | - | tr |
| Iso-C<sub>17:0</sub> | 1.1 | tr | tr |
| C<sub>17:1</sub> ω8*c* | tr | 0.6 | tr |
| C<sub>17:0</sub> | tr | 0.5 | tr |
| C<sub>18:3</sub> ω6*c* (6,9,12) | 2.1 | 3.1 | tr |
| C<sub>18:1</sub> ω9*c* | **10.5** | **21.6** | **30.3** |
| C<sub>18:0</sub> | 1.1 | 2.4 | 2.0 |
| C<sub>19:0</sub> cyclo ω8*c* | - | - | - |
| Summed Feature 2 | 0.5 | tr | 2.9 |
| Summed Feature 3 | **25.8** | **24.1** | **12.1** |
| Summed Feature 8 | 0.6 | 2.8 | - |

\*Summed Feature 2 comprised C<sub>12:0</sub> aldehyde ? , Summed Feature 3 comprised C<sub>16:1</sub> $\omega$ 7*c* and/or C<sub>16:1</sub> $\omega$ 6*c*, Summed Feature 5 comprised C<sub>18:2</sub> $\omega$ 6, 9*c* and/or Ante-C<sub>18:0</sub>, Summed Feature 8 comprised C<sub>18:1</sub> $\omega$ 7*c*.
–, Not detected, less than 0.5% as tr. Fatty acids present at >10% are highlighted in bold. Data are presented to two decimal places.

### Phylogeny analysis

The 16S rRNA sequences of CAAS 2-6^T^ (OQ110573) and CAAS 2-13^T^ (OQ110569) were identical to the Sanger sequencing results. Phylogenetic analysis revealed that strain CAAS 2-6^T^ displayed the highest similarity with *A. gerneri* DSM 14967^T^ (97.7%), followed by *A. kanungonis* PS-1^T^ (97.6%) and *A. tandoii DSM 14970*^T^ (97.5%). In contrast, strain CAAS 2-13^T^ shared 99.6% *16S rRNA* gene sequence similarity with *A. kanungonis* PS-1^T^, with *≤*98.5% similarity to other *Acinetobacter* species. The ML and NJ phylogenetic trees (Fig. S2) delineated the evolutionary relationships between both strains and 96 type strains of validly published *Acinetobacter* species. Strain CAAS 2-6^T^ formed a distinct lineage within the genus, while strain CAAS 2-13^T^ clustered with *A. kanungonis* PS-1^T^ (99% bootstrap support), yet maintained sufficient genetic divergence to suggest novel subspecies status. Notable clade formations included four cohesive clusters: (i) *A. idnjacnsis* Mll^T^, *A. lwoffii* DSMZ 2403^T^ and *A. mesopotamicus* GC2^T^; (ii) *A. pakistanensis* NCCP 644^T^ and *A. movanagherensis* Movanagher 4^T^; (iii) *A. septicus* AKO01^T^ and *A. uusingii* LUH 3792^T^; (iv) *A. dijkshoorniae* JVAP01^T^, *A. geminorum* J00019^T^, *A. calcoaceticus* NCB 22016^T^, *A. pitti* CIP 70.29^T^ and *A. lactucae* NRRL B-41902^T^. MLSA-based ML phylogeny (Fig. **2**) demonstrated strains CAAS 2-6^T^ and CAAS 2-13^T^ occupy independent subclades when compared to 53 *Acinetobacter* species, a pattern corroborated by NJ analysis (Fig. S3). These polyphasic phylogenetic data conclusively establish CAAS 2-6^T^ as a novel species within *Acinetobacter* genus, and CAAS 2-13^T^ as a novel subspecies of *A. kanungonis*.

**FIG 2.**
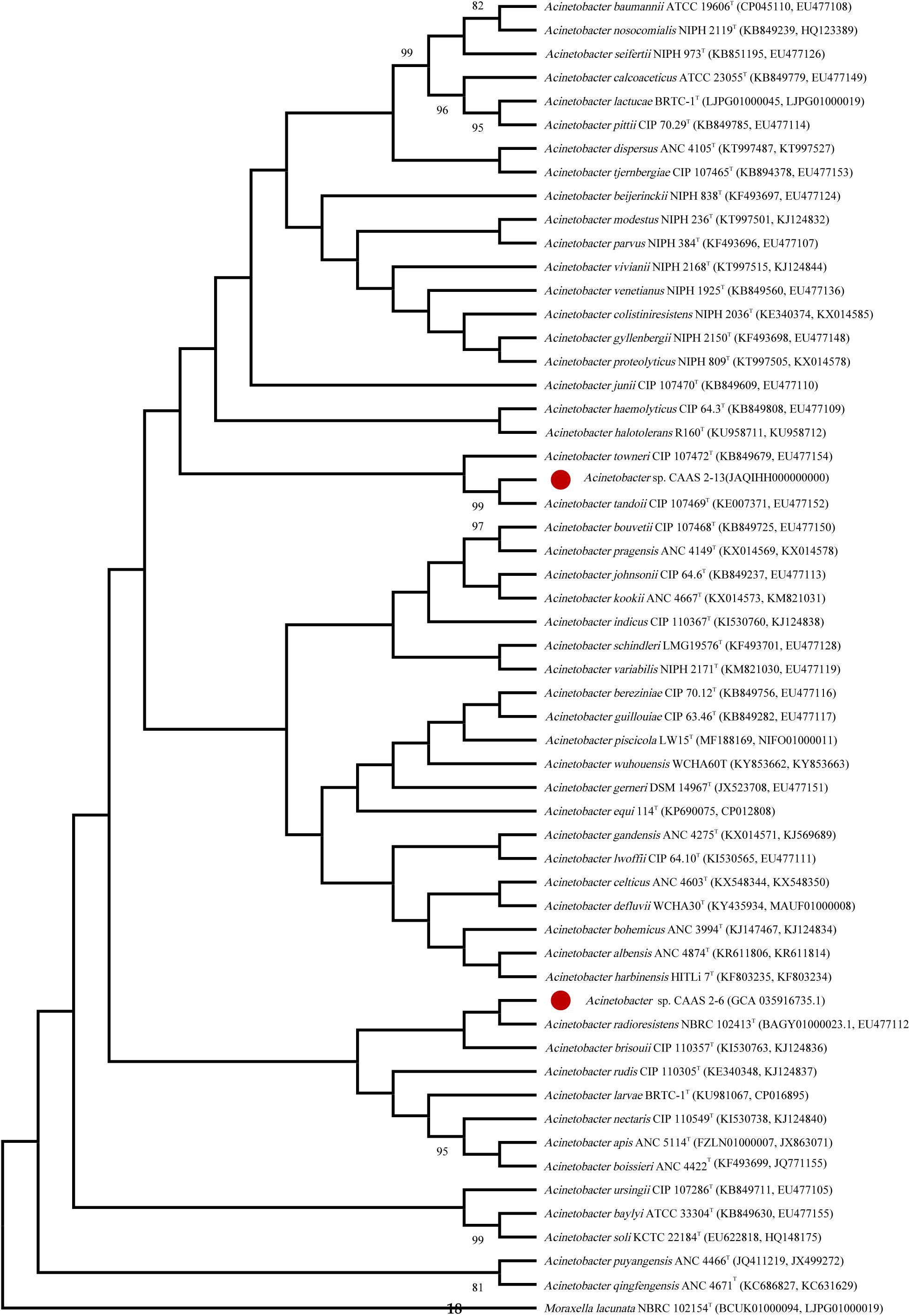
The ML phylogenetic tree inferred from a concatenated alignment of *gyrB* (664 bp) and *rpoB* (742 bp) gene sequences, including strains CAAS 2-6^T^ and CAAS 2-13^T^ alongside 53 type strains of *Acinetobacter* species. *Moraxella lacunata* NBRC 102154^T^ served as the outgroup. Bootstrap support values (*≥*70%) derived from 1,000 replicates are indicated at branch nodes.

### Genomic features and analyses

A total of 1.236 Gb of raw DNA sequences were obtained for strain CAAS 2-6^T^ through sequencing, which were assembled into 73 scaffolds with a coverage of 469*×*. The assembled draft genome of strain CAAS 2-6^T^ was estimated to be at least 2,593,745 bp in size, with a G+C content of 44.4%. Within the assembled draft genome, a total of 2,344 genes, 59 *tRNAs*, 3 *rRNAs*, and 0 CRISPRs were predicted. Functional annotation assigned 2,344, 2,089, 1,783, and 1,803 genes to the NR, eggNOG, KEGG, and Swiss-Prot databases, respectively. The genome sequence of CAAS 2-6^T^ has been deposited in GenBank under accession number JAQIHG000000000/GCA_035916735.1 (Table **2**). For strain CAAS 2-13^T^, 1.113 Gb of raw DNA sequences were obtained, assembled into 48 scaffolds with a coverage of 308*×*. The assembled draft genome of strain CAAS 2-13^T^ was estimated to be at least 3,492,366 bp in size, with a G+C content of 42.0%. A total of 3,332 genes, 66 *tRNAs*, 10 *rRNAs*, and 0 CRISPRs were predicted in the assembled draft genome. Functional annotation assigned 3,273, 2,841, 1,773, and 2,317 genes to the NR, eggNOG, KEGG, and Swiss-Prot databases, respectively. The genome sequence of CAAS 2-13^T^ has been deposited in GenBank under accession number JAQIHH000000000 (Table 2). Circular genome maps of strains CAAS 2-6^T^ and CAAS 2-13^T^, displaying additional features such as open reading frames (*ORFs*), GC content, and GC skew, are presented in Fig. S5.

**TABLE 2.** Genomic assembly results and functional annotation of strains CAAS 2-6^T^ and CAAS 2-13^T^.

| (Property) | CAAS 2-6 | CAAS 2-13 |
| --- | --- | --- |
| Number of contig | 81 | 48 |
| Number of scaffold | 73 | 48 |
| Total sequence length | 2,593,745 | 3,492,366 |
| Coverage (×) | 469 | 308 |
| GC content% | 44.4 | 42.0 |
| ORF number | 2,429 | 3,332 |
| ORF total length | 2,217,186 bp | 3,009,549 bp |
| Longest ORF length | 4,539 bp | 5,142 bp |
| ORF average length | 912.8 bp | 903.2 bp |
| Intergenetic region length | 376,559 bp | 482,817 bp |
| 5S rRNA | 1 | 8 |
| 16S rRNA | 1 | 1 |
| 23S rRNA | 1 | 1 |
| tRNA | 59 | 66 |
| ncRNA | 17 | 18 |
| CRISPRs | 0 | 0 |
| NR annotation | 2,344 | 3,273 |
| eggNOG annotation | 2,089 | 2,841 |
| KEGG annotation | 1,437 | 1,773 |
| Swiss-Prot annotation | 1,783 | 2,317 |
| GO annotation | 1,803 | 2,355 |

Genomic analyses revealed distinct relatedness patterns: CAAS 2-6^T^ showed 19.7-22.7% dDDH, 74.5–77.0% ANI, and 72.0–76.8% ANIb with related strains, with highest similarity to *A. indicus* CIP 110367^T^ (Table S3; Fig. S6a). OrthoANI values (74.3-76.8%) further confirmed its divergence (Fig. S7a). CAAS 2-13^T^ exhibited 66.5% dDDH, 96.0% ANI, and 95.6% ANIb with *A. kanungonis* PS-1^T^ but lower relatedness to others (20.2-28.7% dDDH, 74.1-85.1% ANI, and 73.9–84.6% ANIb) (Table S3; Fig. S6b). OrthoANI analysis showed that CAAS 2-13^T^ had 96.0% similarity with *A. kanungonis* PS-1^T^ and 74.7–85.0% with other strains (Fig. S7b). Meeting established thresholds (70% dDDH/95-96% ANI) (82), CAAS 2-6^T^ represents a novel species and CAAS 2-13^T^ constitutes a novel subspecies.

Comparative synteny analysis (Fig. **3**) identified structural variations: CAAS 2-6^T^ displayed fragmented syntenic blocks indicating genome restructuring, whereas CAAS 2-13^T^ and *A. kanungonis* PS-1^T^ showed conserved collinearity with contiguous syntenic blocks, reflecting evolutionary conservation.

**FIG 3.**
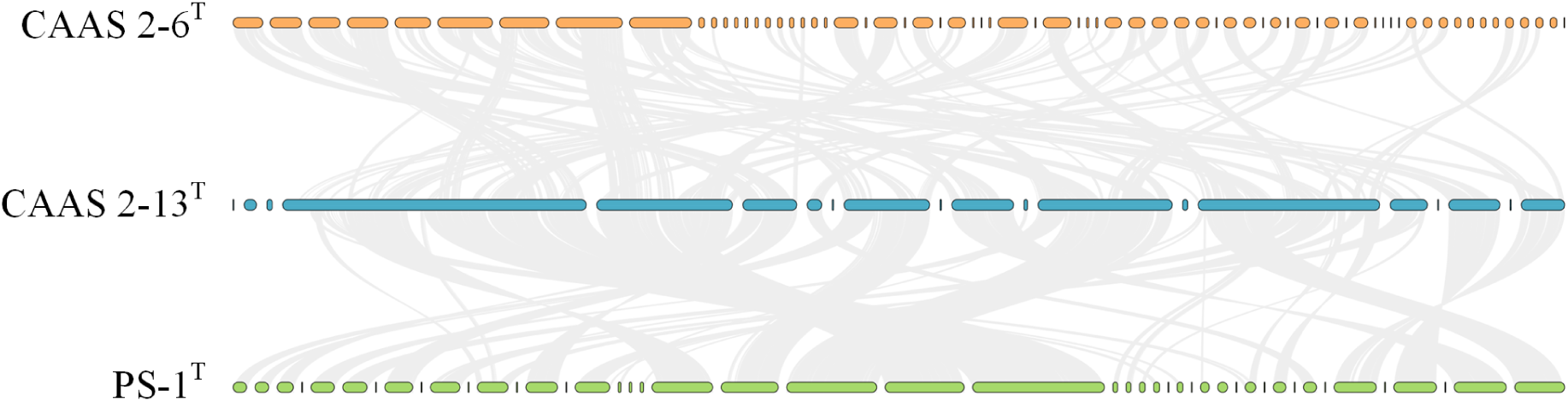
Analysis of collinearity between CAAS 2-6^T^, CAAS 2-13^T^ and *A. kanungonis* PS-1^T^. Orange represents the genome of CAAS 2-6^T^, blue represents the genome of CAAS 2-13^T^, and green represents the genome of *A. kanungonis* PS-1^T^.

### Plastic biodegradation and growth capacity of diverse *Acinetobacter* strains

Type strains *A. gerneri* 9A01^T^ = KCTC 12415^T^ (abbreviated 12415), *A. wuhouensis* WCHA60^T^ = GDMCC 1.1100^T^ (1100), *A. chinensis* WCHAc010005^T^ = GDMCC 1.1232^T^ (1232), *A. sichuanensis* WCHAc060041^T^ = GDMCC 1.1383^T^ (1383) and *A. chengduensis* WCHAc060005^T^ = GDMCC 1.1622^T^ (1622) served as reference controls alongside target isolates CAAS 2-6^T^ and CAAS 2-13^T^ in plastic degradation assays. After 7-day incubation in modified mineral salt medium with single plastic substrates (PLA, PBAT, PET, PBT, PBS, or PBSA), turbidity measurements revealed differential growth capacities (Fig. **4**a). Both CAAS 2-6^T^ and CAAS 2-13^T^ showed no observable growth in PET-amended medium, while *A. sichuanensis* GDMCC 1.1383^T^ exhibited consistently poor growth across all plastics. Significantly enhanced growth was observed in aliphatic polyesters (PLA, PBAT, PBS, PBSA) compared to aromatic polyesters (PET, PBT), aligning with the established recalcitrance of aromatic polyesters to biodegradation. Degradation efficiency ranking indicated: CAAS isolates > four reference strains > *A. sichuanensis* GDMCC 1.1383^T^.

**FIG 4.**
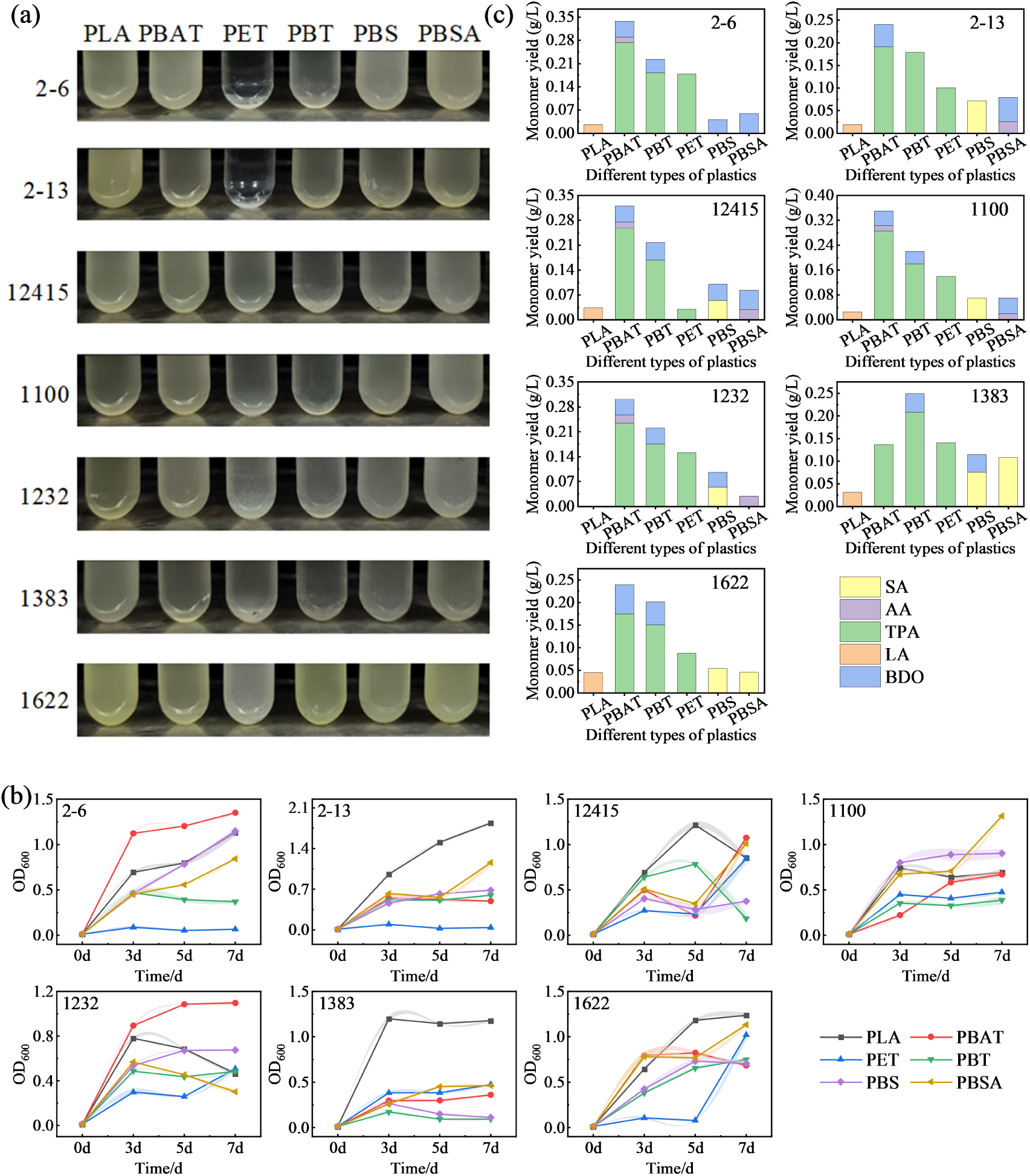
Substrate-dependent growth of *Acinetobacter* strains on synthetic polyesters. (a) Culture phenotypes, (b) Growth kinetics (OD_600_), (c) Monomer quantification by HPLC.

Plastic degradation capacities of seven *Acinetobacter* strains were systematically evaluated by monitoring growth kinetics (OD_600_) and monomer yields across six polyester substrates: PLA, PBAT, PET, PBT, PBS, and PBSA (Fig. **4**b,c). The results demonstrated that strain *CAAS 2-6^T^* exhibited optimal PLA-driven growth (OD_600_=1.13 *±* 0.04) attributed to efficient lactate metabolism, correlating with low lactate accumulation (0.03 g/L). Peak biomass occurred on PBAT (OD_600_=1.35 *±* 0.01), with a terephthalic acid yield of 0.27 g/L. This yield was the highest among terephthalate-containing substrates, in the order of PBAT > PBT > PET. Strain *CAAS 2-13^T^* demonstrated maximal PLA growth (OD_600_=1.84 *±* 0.02) with minimal lactate accumulation (<0.02 g/L). Terephthalic acid production declined progressively: PBAT (0.19 g/L) > PBT (0.18 g/L) > PET (0.10 g/L). PBS degradation yielded succinic acid (0.07 g/L), while PBSA degradation produced both adipic acid (0.03 g/L) and 1,4-butanediol (0.05 g/L). Five reference strains exhibited habitat-independent degradation capacities (OD_600_ range: 0.19-1.31), confirming genus-wide plastic-degrading potential. These findings demonstrate habitat-driven adaptive evolution confers substrate-specific polyester degradation capabilities in *Acinetobacter*: (i) Specialized aliphatic polyester (PLA/PBS/PBSA) degradation in CAAS strains; (ii) Convergent evolutionary adaptation enhancing co-habitat strain efficiency; (iii) Genus-wide ecological versatility enabling synthetic polyester degradation. This work establishes *Acinetobacter* as a prime candidate for enzymatic plastic upcycling, with significant bioremediation potential in environmental biotechnology.

### Expression and enzymatic degradation, and structural analysis of multicopper oxidase AbMCO and alkane hydroxylase AlkB

Based on genomic data, we cloned the coding sequences of multicopper oxidase AbMCO and alkane hydroxylase AlkB for heterologous expression in the pET-28a(+) system. Recombinant proteins from various Acinetobacter strains were successfully expressed in *E. coli* BL21(DE3) under the control of the lac promoter with N-terminal His_6_-tags. Protein samples from recombinant *E. coli* (whole-cell lysates, soluble fractions, and inclusion bodies) were analyzed by SDS-PAGE (12% separating gel). Each recombinant protein migrated as a predominant single band (Fig. S5). Calculated molecular masses excluding the His-tag are approximately 70.8 kDa for 2-6AbMCO1, 66.8 kDa for 2-13AbMCO 1, 69.4 kDa for 2-13AbMCO 2, 66.1 kDa for 12415AbMCO, 67.4 kDa for 1100AbMCO, 65.4 kDa for 1232AbMCO, 63.1 kDa for 1383AbMCO, 66.7 kDa for 1622AbMCO, 46.2 kDa for 2-13alkB, and 46.2 kDa for 1100alkB.

Due to inclusion body formation by recombinant alkane hydroxylase AlkB, refolding was performed with urea according to the HyPur™ T Ni-NTA 6FF Gravity Column protocol (Sangon Biotech, C600791-0010), followed by affinity purification using Ni-NTA resin. Initial purification targeted soluble fractions of 2-6AbMCO1 and 2-13AbMCO2, along with refolded soluble from 2-13AlkB. Purified recombinant enzymes (200 µL) were incubated with 800 µL of reaction buffer and polyester substrates (0.2 g/L powder, <100 µm particle size) at 37°C with agitation (180 rpm) for 120 h. After incubation, the supernatants were analyzed by HPLC. For validation assays, rapid protein purification was conducted at 4°C, with enzymatic depolymerization initiated within 24 h. HPLC analysis (Fig. **5**) revealed higher monomer yields from AlkB-mediated degradation versus AbMCO, particularly for terephthalic acid from PBAT (>0.9 g/L; Fig. **5**a). Among all AbMCO variants, 2-6AbMCO1 demonstrated optimal catalytic performance: total product yield of 3.8 g/L with enzymatic productivity of 9.4 g product/g enzyme. In contrast, 12415AbMCO showed significantly reduced efficiency (Fig. **5**b,c).

**FIG 5.**
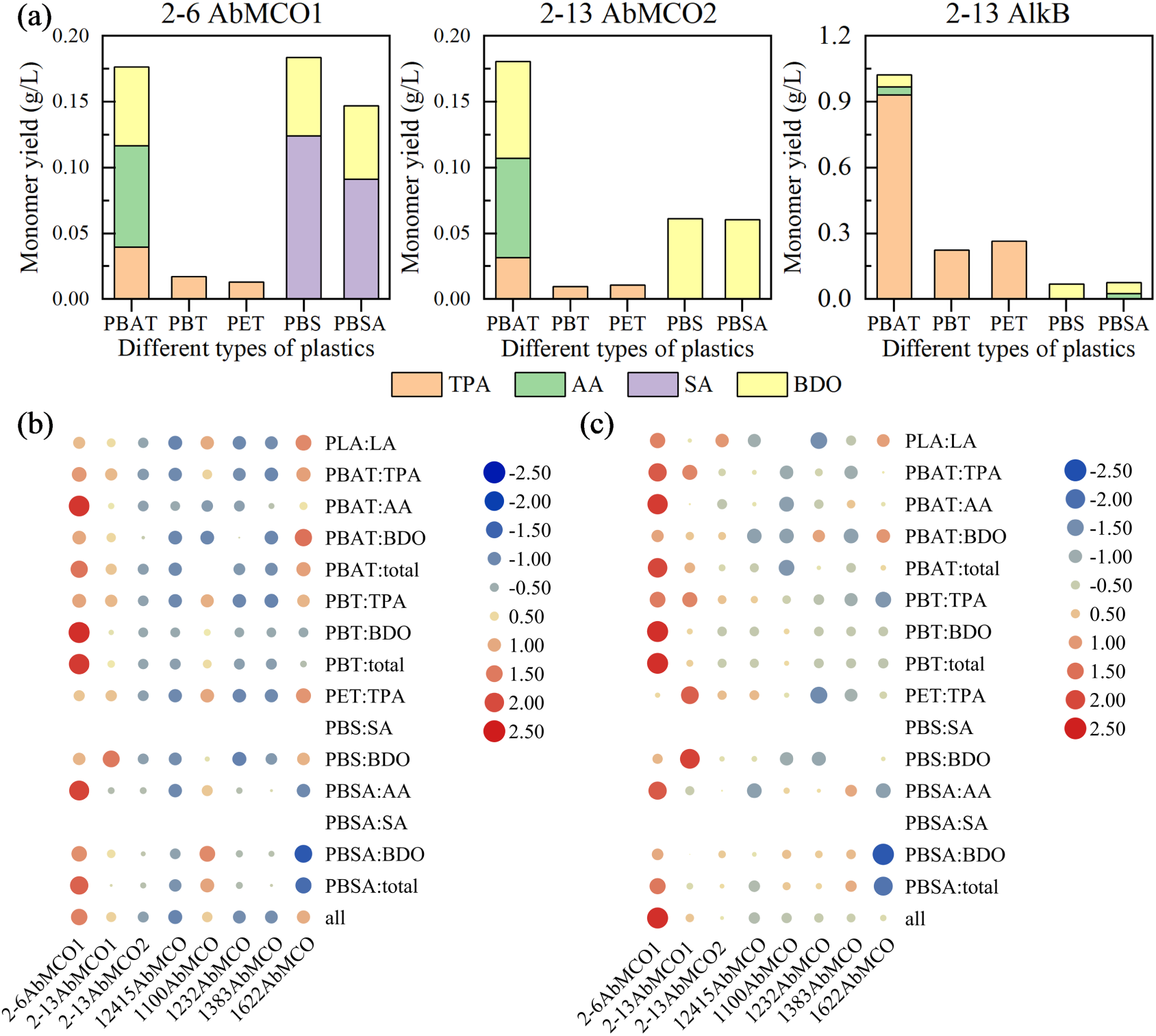
HPLC analysis of monomeric products from plastic degradation by heterologously expressed AbMCO and AlkB. (a) Primary enzymatic depolymerization assay, (b) Product concentration (g/L) in validation assay, (c) Enzymatic yield (g/g) in validation assay. Abbreviations: SA, succinic acid; AA, adipic acid; TPA, terephthalic acid; LA, lactic acid; BDO, 1,4-butanediol.

Multiple sequence alignment of multicopper oxidase AbMCO and alkane hydroxylase AlkB (Fig. S6) revealed divergent conservation patterns. AbMCO exhibited poor conservation in the mid-rear and C-terminal regions, correlating with predicted random coil structures. The multicopper oxidase AbMCO, which is highly conserved across *A. baumannii* strains, exhibits *in vitro* enzymatic activity only upon supplementation with *≥*1 µM copper ions, demonstrating its obligate copper dependence (83). AlkB showed low sequence conservation at N-terminal, mid-rear, and C-terminal segments, while conserved domains predominantly adopted α-helical conformations. Despite significant sequence diversity, AlkB enzymes consistently catalyze the initial hydroxylation of hydrocarbons, with conserved structural features in medium-length alkane biodegradation (84).

Phylogenetic analysis placed 12415AbMCO and 2-13AbMCO1 on basal branches with significant genetic divergence. A highly supported clade (Bootstrap=97-100) comprising 1100AbMCO, 1232AbMCO, 1383AbMCO and 1622AbMCO displayed short branch lengths, indicating strong sequence conservation. 2-13AlkB and 1622AlkB occupied distal positions, while the ancestral node of 12415AlkB-1383AlkB showed moderate support (Bootstrap=64-87), warranting further validation. AlphaFold structural comparisons yielded low RMSD values among 1100AbMCO, 1232AbMCO, 1383AbMCO and 1622AbMCO, confirming high structural similarity. Despite shared ecological origins, 2-6AbMCO1 diverged structurally from 2-13AbMCO1/2-13AbMCO2. 1383AlkB showed marked structural deviation from other AlkBs, even though 2-13AlkB was phylogenetically distant (Fig. **6**).

**FIG 6.**
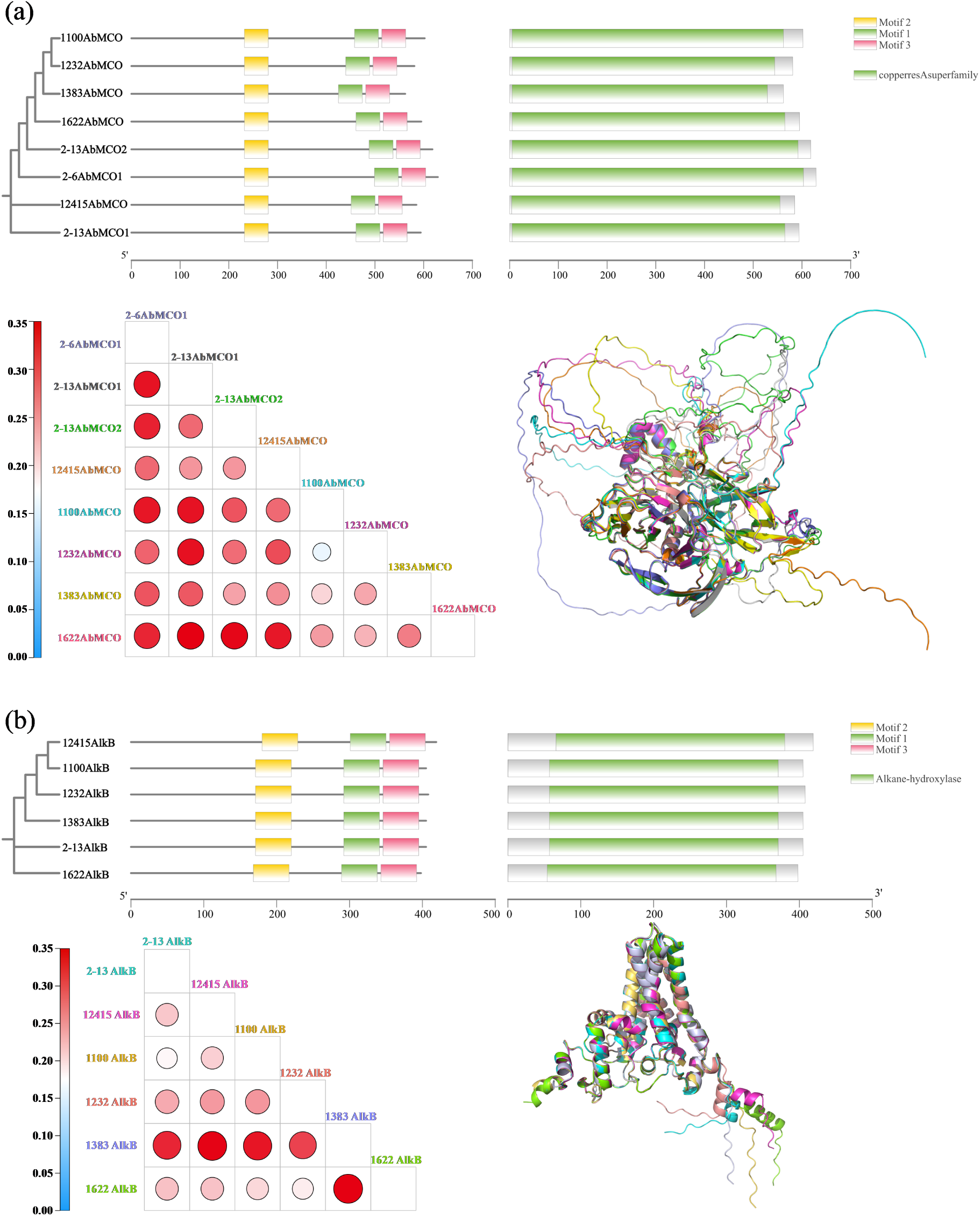
Comparative analysis of evolutionary relationships and structural features among recombinant AbMCO (a) and AlkB (b) from various Acinetobacter strains. Top: Phylogenetic tree based on amino acid sequences. Bottom left: The root-mean-square deviation (RMSD) values (in Å) of the aligned AlphaFold-predicted model structures, indicating the similarity between protein structures. Bottom right: Visual representations of the structural alignment results. The color of protein labels in the phylogenetic tree corresponds to the color of the same proteins in the structural comparison visualizations.

Structural alignments of AlphaFold-predicted AbMCO models revealed a predominantly β-sheet-rich fold, characterized by minimal α-helical content and extensive random coil regions. A disordered segment within this structure correlates with low prediction confidence, likely attributable to limited homology with experimentally resolved templates. Notably, its substrate-binding domain forms a negatively charged catalytic pocket, where hydrophobic residues line the binding site and substrate access is regulated by rigid α-helical gatekeeping elements. Conversely, AlkB adopts a scaffold-like architecture typical of non-heme diiron monooxygenases. This structure comprises: (i) Six transmembrane helices (TM1–TM6) constituting a substrate-access channel, (ii) An N-terminal cytosolic catalytic domain housing a hydrophilic di-iron center essential for oxygen activation, and (iii) A membrane-embedded C-terminus adjacent to the catalytic domain that stabilizes the complex (Fig. **6**).

Based on plastic depolymerization activity and AlphaFold structural alignments, molecular docking of representative enzymes 2-6AbMCO1 and 2-13AlkB with 12 oligomeric substrates (ABT, BAB, BABTB, BSB, BT, BTB, BTBT, LLLL, MHET, MHET4, TBT, TBTBT) revealed distinct binding topologies: 2-6AbMCO1 accommodated substrates at five binding sites—Site 1 (BSB/BTB/LLLL/MHET), Site 2 (ABT/TBT/BAB/BT), Site 3 (BABTB/TBTBT), and exclusive Sites 4-5 for BTBT and MHET4 respectively; whereas 2-13alkB utilized seven loci predominantly at its base—including a shared catalytic pocket (Site 1: ABT/BAB/BT/BTB/BTBT/LLLL), vicinal sites for MHET4/BSB/TBT/BABTB (Sites 2-5), a waist-positioned site for TBTBT (Site 6), and an apical site for MHET (Site 7)—notably exhibiting differential residue lining patterns within the common pocket occupied by six substrates (Fig. **7**).

**FIG 7.**
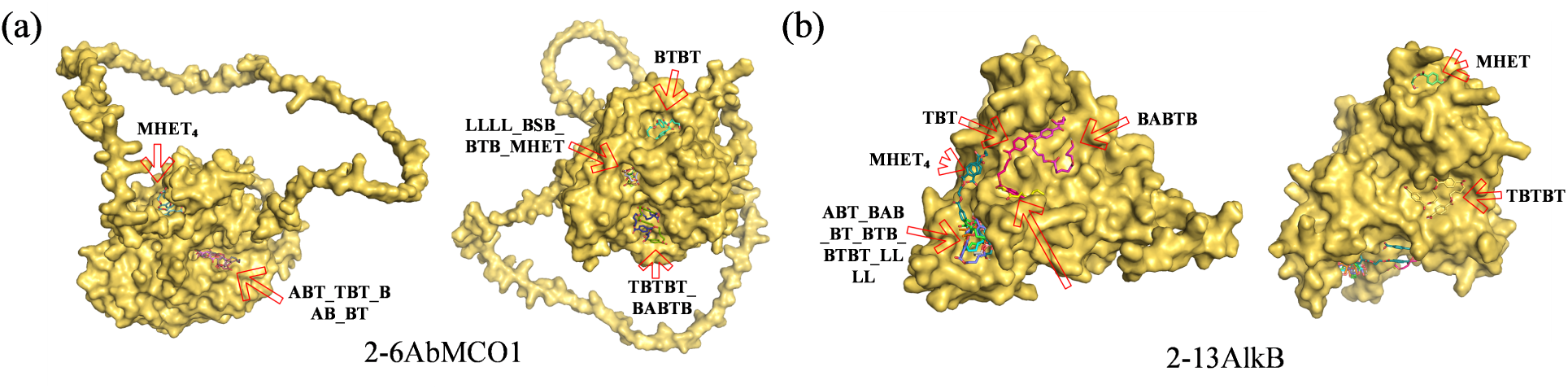
Molecular docking of 2-6AbMCO1 and 2-13AlkB. (a) 2-6AbMCO1 binding sites. (b) 2-13alkB binding sites.

## CONCLUSION

In this study, strains CAAS 2-6^T^ and CAAS 2-13^T^ were isolated from landfill leachate and a strawberry cultivation field, respectively. These novel plastic-degrading strains were taxonomically identified as *Acinetobacter lentus* sp. nov.and *Acinetobacter kanungonis* subsp. *fragariae* subsp. nov.. Both CAAS 2-6^T^ and CAAS 2-13^T^ alongside five reference strains were demonstrated the ability to degrade various plastic types. Notably, strains CAAS 2-6^T^ and CAAS 2-13^T^ grew on plastics (PLA, PBAT, PBT, PBS, PBSA) as the sole carbon source, reaching an OD_600_ of 0.3 within 3–7 days. Genomic mining revealed the expression of multicopper oxidase AbMCO and alkane hydroxylase AlkB, with enzymatic depolymerization assays confirming their involvement in plastic degradation. *Acinetobacter* Species exhibit broad biotechnological potential for degrading recalcitrant pollutants (e.g., plastics, hydrocarbons, pesticides, and plasticizers), underscoring their promise for bioremediation applications. Future studies should characterize additional plastic-degrading enzymes from strains CAAS 2-6^T^ and CAAS 2-13^T^ to elucidate their metabolic pathways.

### Description of *Acinetobacter lentus* sp. nov

*Acinetobacter lentus* (len’tus. L. masc. adj. lentus, slow, delayed, referring to slow growth).

The strain CAAS 2-6^T^, isolated from landfill leachate, forms circular, regular-edged, smooth colonies of 0.5–1.0 mm in diameter on LB plates after being incubated at 30°C for 2 days. The cells are rod-shaped, measuring 0.6–0.7 µm *×* 0.7–1.3 µmm. The strain grows within a temperature range of 15–40°C, with an optimum growth temperature of 25–40°C. It can grow in the presence of 0–6.0% (w/v) NaCl, with optimal growth observed at 0–4.0% (w/v) NaCl. The pH range for growth is 6.0–8.0, with an optimum pH of 7.0–7.5. *16S rRNA* sequence (GenBank accession number: OQ110573) analysis revealed a similarity of 97.7%, 97.6%, 97.5%, and 97.5% with *A. gerneri* DSM 14967^T^, *A. kanungonis* PS-1^T^, *A. tandoii* DSM 14970^T^ and *A. junii* CIP 64.5^T^, respectively. Evolutionary tree analysis based on *16S rRNA* sequences, as well as the housekeeping genes *gyrB* and *rpoB*, showed that CAAS 2-6^T^ formed a separate cluster. The strain is negative for oxidase, gelatinase, and nitrate reduction, and cannot assimilate adipic acid, malic acid, and sodium citrate. It cannot utilize L-arabinose, D-xylose, galactose, and amygdalin. However, it tests positive for the Voges-Proskauer reaction, can assimilate D-glucose, and can ferment D-glucose, D-mannitol, inositol, sorbitol, rhamnose, sucrose, melibiose, amygdalin, and L-arabinose to produce acid. It can also utilize glucose, fructose, mannose, N-acetyl-glucosamine, amygdalin, arbutin, esculin, maltose, sucrose, and starch. The G+C content of the genome (GenBank accession number: JAQIHG000000000/GCA_035916735.1) is 42.0 mol%. The highest dDDH and ANI values were observed with *A. indicus* CIP 110367^T^, at 22.7% and 77.0%, respectively. The main fatty acids are Summed Feature 3 (comprising C_16:1_ *ω*7*c* and/or C_16:1_ *ω*6*c*, 25.8%), C_16:0_ (25.7%), and C_18:1_ *ω*9*c* (10.5%). The major polar lipids are diphosphatidylglycerol (DPG), phosphatidylethanolamine (PE) and phosphatidylglycerol (PG). Based on the experimental results of physiological and biochemical characteristics, genomic homology, and content, the CAAS 2-6^T^ strain is considered a new species of the genus *Acinetobacter*. Due to its distinguishing growth rate compared to other species within the genus, it has been named *Acinetobacter lentus* sp. nov. The CAAS 2-6^T^ strain, as the type strain of *Acinetobacter lentus*, has been preserved at the China General Microbiological Culture Collection Center, the Japan Collection of Microorganisms, and the Korean Collection for Type Cultures with preservation numbers GDMCC 1.3951, JCM 36321, and KCTC 8156, respectively.

### Description of Acinetobacter kanungonis subsp. fragariae subsp nov

*Acinetobacter kanungonis* subsp. *fragariae* (frag’ar.i.ae. N.L. fem. n. Fragaria, generic name of strawberry; N.L. gen. fem. n. fragariae, of Fragaria).

The strain CAAS 2-13^T^, isolated from a strawberry field, forms circular, regular-edged, smooth, and flat colonies of 1.5–2.0 mm in diameter on LB plates after being incubated at 30°C for 2 days. The cells are short rod-shaped, with dimensions of 0.5–0.6 µm *×* 0.7–1.2 µm. This strain grows within a temperature range of 15–40°C, with an optimum growth temperature of 20–37°C. It can grow in the presence of 0–5.0% (w/v) NaCl, with optimal growth observed at 0–3.0% (w/v) NaCl. The pH range for growth is 4.5–8.0, with the most suitable pH being 5.0–7.5. Analysis of the *16S rRNA* sequence (GenBank accession number: OQ110569) revealed a high similarity of 99.6% to the *16S rRNA* sequence of *A. kanungonis* PS-1^T^. The *16S rRNA* sequence-based phylogenetic tree clustering is closest to *A. kanungonis* PS-1^T^, while the phylogenetic clustering based on the housekeeping genes *gyrB* and *rpoB* is closest to *A. tandoii* ClP 1074697^T^. The strain is negative for oxidase, Voges-Proskauer reaction, and nitrate reduction. It cannot ferment D-glucose, inositol, sorbitol, rhamnose, melibiose, amygdalin, or L-arabinose to produce acid, and it cannot assimilate D-glucose or adipic acid. Additionally, it cannot utilize L-arabinose, ribose, D-xylose, galactose, mannose, amygdalin, arbutin, sucrose, or starch. However, it is gelatinase-positive, can assimilate malic acid and sodium citrate, and can utilize glucose, fructose, N-acetyl-glucosamine, esculin, and maltose. The G+C content of the genome (GenBank accession number: JAQIHH000000000) is 42.0 mol%. The highest values of dDDH and ANI with *A. kanungonis* PS-1^T^ are 66.5% and 96.0%, respectively. The major fatty acids are C_16:0_ (25.5%), Summed Feature 3 (comprising C_16:1_ *ω*7*c* and/or C_16:1_ *ω*6*c*, (24.1%), and C_18:1_ *ω*9*c* (21.6%). Based on a comprehensive evaluation of physiological and biochemical characteristics, genomic homology, and content, despite the similarity and average nucleotide identity of the *16S rRNA* sequence being above the recommended thresholds, the dDDH is slightly below the recommended cutoff value. Furthermore, there is a certain evolutionary distance between TYGS and *A. kanungonis* PS-1^T^ in the genomic phylogenetic tree. Therefore, the strain CAAS 2-13^T^ is identified as a novel subspecies of *A. kanungonis* PS-1^T^. Given its distinct isolation source from *A. kanungonis* PS-1^T^, it is named *Acinetobacter kanungonis* subsp. *fragariae* subsp. nov. The strain CAAS 2-13^T^, representing *Acinetobacter kanungonis* subsp. *fragariae*, has been preserved at the China General Microbiological Culture Collection Center, the Japan Collection of Microorganisms, and the Korean Collection for Type Cultures with preservation numbers GDMCC 1.3956, JCM 36322, and KCTC 8157, respectively.

## ACKNOWLEDGMENTS

We sincerely appreciate the help of Professor Ma Shi-Chun from Biogas Institute of Ministry of Agriculture and Rural Affairs in fatty acids composition analysis using the MIDI system, and the China Agricultural Culture Collection Center (ACCC) for providing the polar lipid profile report. This work was financially supported by National Key R&D Program of China (Grant No. 2023YFE0104900-04-04), Agricultural science and Technology Innovation Project of Chinese Academy of Agricultural Sciences (CAAS-ASTIP-2016-BIOMA), Qinghai Science and Technology program (2022-NK-124) and Chengdu Science and Technology Program (NASC2023TD09).

## FUNDING

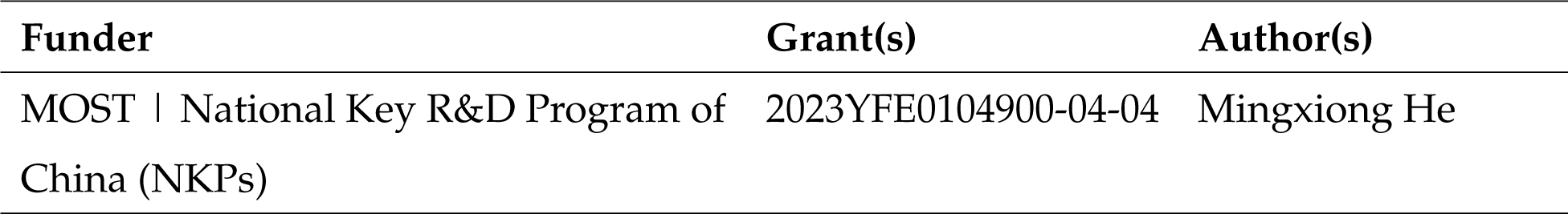

## AUTHOR CONTRIBUTIONS

Mengli Xia, Conceptualization, Formal analysis, writing - original draft | Yuandong Zhao, Formal analysis, Investigation | Yiran Ma, Resources, Validation | Bo Wu, Formal analysis, Visualization, Writing – original draft | Guoquan Hu, Investigation, Methodology | Yanwei Wang, Project administration, Methodology, Supervision | Mingxiong He, Funding Acquisition, Supervision, Writing – review and editing..

## DATA AVAILABILITY STATEMENT

The GenBank/EMBL/DDBJ accession number for the genome and *16S rRNA* gene sequences of strain *Acinetobacter lentus* CAAS 2-6^T^ are JAQIHG000000000/GCA_035916735.1 and OQ110573 respectively. The GenBank/EMBL/DDBJ accession number for the genome and *16S rRNA* gene sequences of strain *Acinetobacter kanungonis* subsp. *fragariae* CAAS 2-13^T^ are JAQIHH000000000 and OQ110569 respectively.

## CONFLICTS OF INTEREST

The authors declare no conflict of interest.

## ADDITIONAL FILES

The following material is available online.

## Supplemental material

Supplemental material (AEM00000-00-s0001.docx). Figures S1 to S11 Tables S1 to S3.

